# The extracellular matrix dictates ovarian cancer cell migration in an in vivo-derived circulating environment

**DOI:** 10.1101/2025.10.01.679803

**Authors:** Zixu Wang, Alphonse Boché, Laure Chardin, Franck Carreiras, Marie-Claire Schanne-Klein, Yong Chen, Alexandra Leary, Ambroise Lambert, Carole Aimé

## Abstract

Ovarian cancer (OC) disseminates via ascites and interaction with peritoneal extracellular matrix (ECM). To dissect the crosstalk between ECM and ascites in OC cell migration, we developed an ovarian tumor-on-chip integrating OC tumor spheroids, perfusion-induced shear stress of fluid supplemented with key components of ascites or patient-derived ascites, and biomimetic ECMs mimicking either early-stage basement membrane or late-stage connective tissues. We evaluated the individual and combined effects of two key ascitic components, fibronectin and TGF-β, and compared these results with the perfusion of patient ascites. Results showed that cell migration, morphology, and epithelial-to-mesenchymal transition (EMT) markers such as the reorganization of vimentin cytoskeleton depend strongly on ECM composition, regardless of biochemical cues. Fibronectin and TGF-β synergistically enhanced migration and EMT signatures, especially on basement membrane-rich ECM. Patient ascites further promoted migration but did not override ECM-driven migration patterns. Our findings show that clinical ascites perfusion exemplifies the ECM-dependence of cancer cell migration. This highlights that ECM protein composition is a dominant regulator of OC cell migration, providing key insights for *in vitro* tumor modeling and therapeutic strategies targeting the metastatic microenvironment.

## 1. Introduction

The poor prognosis of ovarian cancer (OC) is attributed to diagnosis at an advanced stage with diffuse spread to the surface of the peritoneal cavity [1]. Many studies have demonstrated a correlation between the migratory capacity of OC cells and the rigidity of the underlying substrate [2,3]. However, physical factors that affect cell migration behavior also include ECM fiber morphology [4] and composition [5,6], mechanical stimulation including shear stress [7] and tensile stress [8]. Ascites is a pathological fluid in the peritoneal cavity, which builds up in the course of ovarian cancer (OC) [9]. Malignant ascitic fluid, containing abundant tumor-promoting cytokines, chemokines, growth factors, and proteinases, provides a distinct type of tumor microenvironment, playing a critical role in OC metastasis [10,11]. There is increasing evidence that ascites contributes to the promotion of epithelial-to-mesenchymal transition (EMT), shifting tumor cells towards a stem cell-like phenotype [12-14]. This can be associated with the expression switch of vimentin, a mesenchymal intermediate filament protein, whose upregulation is a hallmark of EMT phenotypic shift and a functional mediator of mesenchymal properties like motility and invasion.

Transforming growth factor-beta 1 (hereafter abbreviated as TGF-β) is a significant tumor-promoting cytokine found in the malignant ascites of OC patients [15]. Primarily secreted by tumor-associated macrophages, TGF-β in ascites enhances cancer cell migration, adhesion, and chemoresistance, thus decreasing survival [16-18]. Elevated levels of TGF-β in ascites are indicative of early recurrence and a poor prognosis [19,20]. TGF-β has been found to be crucial in both EMT and the reverse mesenchymal-to-epithelial transition (MET), playing a key regulatory role in spheroid formation and implantation [21,22]. Additionally, TGF-β in ascites stimulates the proliferation, activation, and deposition of type I collagen by omental fibroblasts, and enhances the responsiveness of ovarian tumor cells to other growth factors present in ascites, such as heparin-binding epidermal growth factor [23,24]. On the other hand, elevated levels of fibronectin have been observed in the stroma of ovarian tumors and in the ascites fluid of patients with OC [25,26]. A significant correlation exists between increased fibronectin expression, advanced tumor stages, and decreased overall survival in OC [27,28]. Functional studies have shown that fibronectin can drive the proliferation of ovarian tumor cells and facilitate metastasis by modulating the cell adhesion and invasion capabilities [29].

Investigating the impact of ascites requires *in vitro* models integrating key components of the cell microenvironment. With this objective, we have elaborated an ovarian tumor-on-chip (OToC) integrating 1/ ECM models recapitulating the early and late stages of OC, 2/ OC spheroids, 3/ controlled shear stress simulating the one in the peritoneal cavity, and 4/ perfusion of key ascites components, fibronectin and TGF-β, isolated or combined. Ultimately, we converge this model ascites approach with a more clinical approach by perfusing patient-derived ascites. By doing so, we assess the crosstalk between ECM, OC spheroids and the OC-specific dynamic circulating microenvironment. This work indicates that the perfusion of fibronectin or/and TGF-β, while acting synergistically, enhances spheroid migration, with cell migration showing a strong dependence on the ECM composition. This was also confirmed when perfusing withpatient-derived ascites, further underscoring the importance of incorporating ECM complexity in *in vitro* models to reproduce metastatic processes. Moreover, this work highlights the need for the development of tumor-on-chip for rationally dissecting tumor–microenvironment interactions and supports the development of ECM-targeted therapeutic strategies.

## 2. Methods

### 2.1. Fabrication of the porous supports

Porous supports similar to the ones that are commercially available from the Mesobiotech® company were fabricated by photolithography and soft-lithography in the lab as previously reported (Supporting Information 1.1.) [5]. The width of the honeycombs was set at 400 μm, with a frame width of 50 μm and a thickness of 50 μm.

### 2.2. ECM fabrication

Type I collagen was extracted and purified from rat tail tendons as previously described, except that we used 3 mM hydrochloric acid instead of 500 mM acetic acid [30,31]. Collagen purity was assessed by electrophoresis and its concentration estimated by hydroxyproline titration [32]. All other chemicals were purchased and used as received. Water was purified with a Direct-Q system (Millipore Co.). Two ECM models were prepared. The first one is an artificial connective tissue (ACT) from type I collagen only. A PBS solution at pH 9 was prepared by slowly adding 1M NaOH to 1X PBS (pH = 7.4) to induce collagen I fibrillogenesis from acidic collagen I solution (1 mg/mL in PBS). In the second model, called artificial basement membrane (ABM), type I collagen (1 mg/mL in PBS), type IV collagen (0.5 mg/mL in PBS, from human placenta) and laminin (10 μg/mL in dionized (DI) water, from Engelbreth-Holm-Swarm murine sarcoma basement membrane) were drop cast by pouring 20 μL of the solution on the patch and dried in air at room temperature in between each step of drop casting in the layer-by-layer process (Figure 1A). Finally, the ECM was washed with DI water, dry in air at room temperature and could be stored in the fridge at 4?.

**Figure 1.**
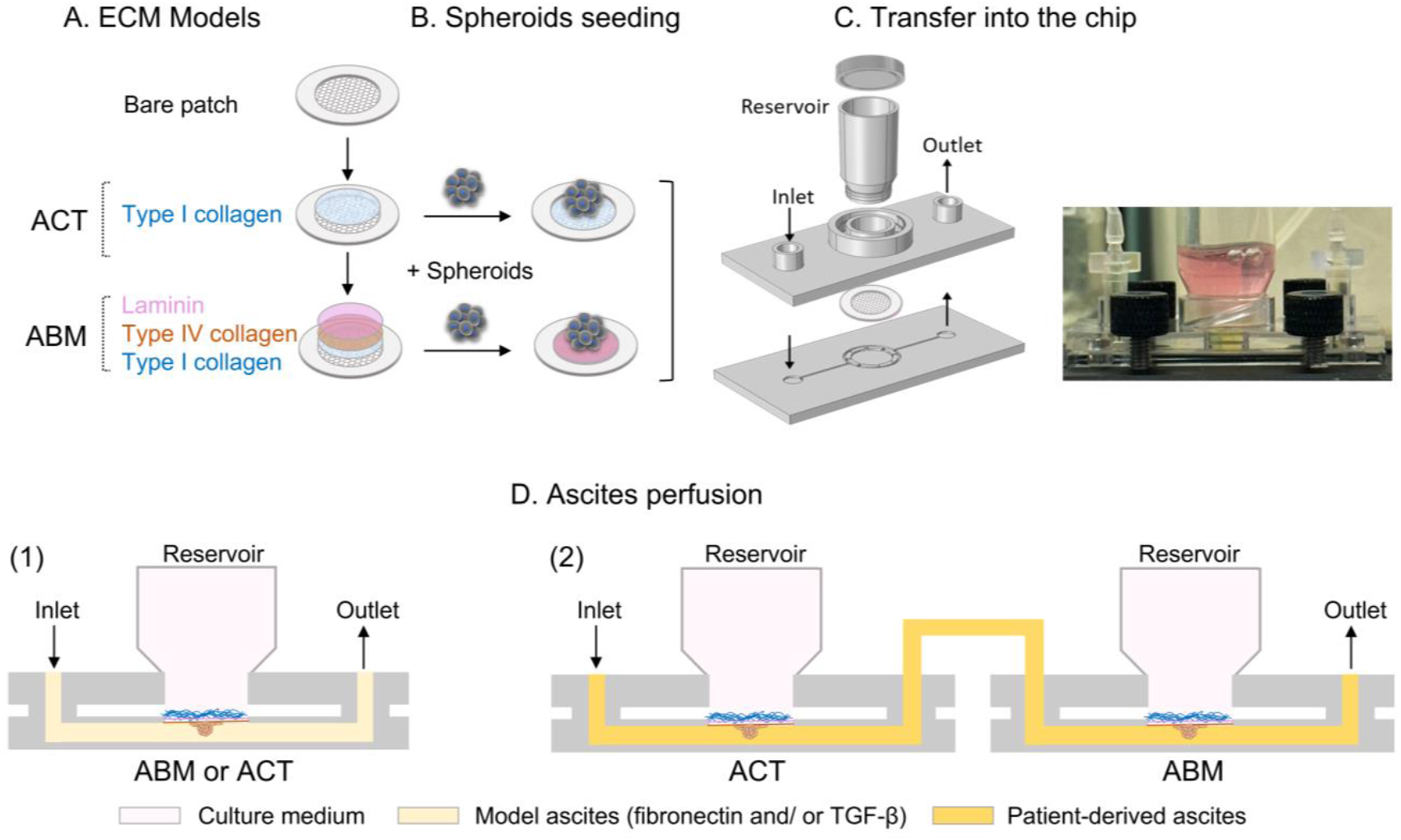
Schematics of the OToC. (A) ECM models: artificial connective tissues (ACT) and artificial basement membrane (ABM), (B) spheroids seeding, and (C) integration of the seeded ECM into the microfluidic chip. (D) Schematics of the microfluidics setup for perfusing (1) model ascites or (2) patient ascites.

### 2.3. Cell Culture

Human ovarian adenocarcinoma SKOV-3 cells (ATCC1, HTB77™) were purchased from ATCC (American Type Culture Collection, Manassas, VA). Cells were cultured in RPMI-1640 glutaMAX containing 0.07 % (v/v) sodium bicarbonate supplemented with 10% fetal bovine serum and 1% (v/v) penicillin streptomycin (all reagents were purchased from Thermo Fisher Scientific). Cells were cultured in T25 cell culture flasks in a humidified air atmosphere with 5% CO_2_ at 37°C.

### 2.4. Spheroid growth

Ovarian tumor spheroid (OTS) growth was performed with a high control over spheroid size (Supporting Information 1.2.) [33]. Briefly, gelatin nanofibers were electrospun on caged-patch having a wall thickness of 200 μm. Then, agarose was coated on the gelatin fibers. After maintaining the spherical patch in the bottom of a 12-well plate using a PDMS ring, SKOV3 cells (4×10^5^ cells/patch) were deposited on the caged-patch. After 3 days culture, uniform spheroids were obtained with a diameter of *ca*. 200 μm. Ovarian tumor cells in spheroids were stained with epithelial (EpCAM and E-cadherin) and mesenchymal (Vimentin) markers (Figure S1-S2).

### 2.5. Seeding ovarian tumor spheroids and integration in the microfluidic chip

Cell migration experiments were carried out on ACT and ABM under perfusion. First, all ECM models were sterilized by immersion in 70% ethanol for 5 min before washing in sterilized PBS 3 times for 5 min each time. The ACT/ABM supports were maintained at the bottom of the dish with a PDMS ring. Four OTS with a uniform diameter of *ca*. 200 μm were selected out using a micropipette and seeded one by one at similar positions on both the ACT and ABM patches (Figure 1B). The number of spheroids seeded is chosen to improve the statistical quality of our data and to ensure that the spheroids are sufficiently separated not to interfere with each other. After incubation on the ECM models for 2 hours at 37? and 5% CO_2_ to ensure firm adhesion of the spheroids to the patches, the patches were loaded into the microfluidic chip (Figure 1C).

### 2.6. Integration in the microfluidic chip

The seeded ECM was then transferred in a commercial microfluidic device (Mesobiotech, France) composed of two plastic plates and a circular culture chamber (∅ 10 mm, h 1.5 mm) at the center of the chip. The chip is closed with magnetic bounding and a mechanical clamper with four hand screws to prevent leakage for cell culture under flow for two days. The upper plate was designed to have two Luer connectors (one inlet and one outlet) and a central reservoir. The inlet channel is divided into six smaller channels to spill the flow into the central cavity while reducing the shear stress. To mimic the shear stress of the peritoneal cavity, an isotropic flow with a shear smaller than a dyn.cm^-2^ was applied in the microfluidic chip. We developed a long-term circulation system, which enables unidirectional flow of the culture medium in the microfluidic chip system without intervention over 48 hours culture time (Figure S3).

The distribution of the shear rate (ν in s^-1^) in the central cavity was simulated by computer fluid dynamics modeling at a flow rate of 20 μL.min^-1^ and calculated to be ν ϵ (0.2-0.4 s^-1^) (Figure S4). Accordingly,

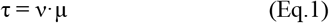

where τ is the shear stress and μ is the dynamic viscosity of culture medium. Thus, τ ϵ (2.4×10^-3^-4.8×10^-3^ dyn.cm^-2^), with a dynamic viscosity of 1.2×10^-2^ dyn.s.cm^-2^. For the given channel with a section area in the central chamber of ∼0.2×10 mm^2^, a linear fluid velocity of ∼0.2 mm.s^-1^ can be obtained, which also give a shear rate in the order of the simulated values.

### 2.7. Identification of patient and collection of samples

All patients provided written informed consent authorizing the use of biological samples obtained during their routine diagnosis and treatment as part of the prospective academic research study OvBIOMARK (NCT03010124). Fresh ascites were obtained from patients with confirmed epithelial OC during primary or secondary laparoscopy. At that time, 15 mL of freshly collected ascites were double centrifuged at 1000 g and 14 000 g within an hour of collection and the supernatant was frozen at -80 °C.

### 2.8. Immunofluorescence and confocal imaging

After 2 days of culture, cells were fixed in 4% paraformaldehyde (PFA) in PBS for 10 minutes, rinsed three times with PBS for 5 min each time. The cells were permeabilized with 0.5% Triton X-100 in PBS for 10 minutes, washed again 3 times in PBS and saturated with PBS containing 3% BSA (with 0.1% Tween 20 and 0.1% sodium azide) for 30 min and washed again. Cells were then incubated overnight at 4°C with Alexa Fluor 543-conjugated vimentin antibody (ab202504, Abcam®) at a 1/1000 dilution and washed 3 times with PBS for 5 min each time to remove unbound vimentin conjugate. After washing, the actin cytoskeleton was stained with a 50 mg/mL Alexa Fluor 488 Phalloidin in PBS (containing 1% DMSO from the original stock solution, Abcam®) for 40 min at room temperature in a dark chamber. Cells were washed 3 times with PBS for 5 min each time to remove unbound phalloidin conjugate. Nuclear DNA was then stained with DAPI (4,6-diamidino-2-phenylindole dihydrochloride, Molecular Probe®) for 15 min at room temperature and washed again. Immunofluorescent labelling was observed with a confocal microscope (LSM710, Zeiss) equipped with 405 (DAPI), 488 (phalloidin), and 543 (anti-vimentin antibody) nm lasers and with LSM ZEN 2009 software. We used 1 μm z-stack intervals and sequential scanning for each wavelength.

### 2.9. Image processing and statistical analysis

The migratory behavior (migrating distance, number of migrating cells and core area, Figure S6) was analyzed using Image J [34]. The staining intensity of vimentin in the core and leading areas were analyzed using ImageJ by measuring the ratio of the integrated fluorescence intensity of a region of interest to the area of this region. More than 4000 cells were analyzed. First, individual cell and nucleus have been labeled using napari-cellpose software [35]. Then, each label enters a FIJI macro designed to extract features and morphologies from each individual cell and nucleus label [36]. Every group of experiments has been run in triplicate and at least 9 tumor spheroids were statistically analyzed for each condition using the Two-Sample t-Test. Finally, the morphology of cells and nuclei were analyzed by principal component analysis (PCA) using the CellTool software [37,38]. PCA allows the determination of the standard deviation (s.d.) that represents the squareroot of the total variance calculated from all the measured morphological features, and hence quantifies the cellular heterogeneity. PCA analysis has been run with four spheroids per patch, and as triplicates of patch for each condition. The number of isolated cells in the leading area analyzed per condition is reported in Table S1.

## 3. Results

We developed ECM models composed of only ECM proteins drop cast on a porous support. This support with 400 μm-wide pores aims at enabling the handling of mostly unsupported tissues, thus reaching physiological stiffness, and getting rid of the mechanical contribution of the support. In order to mimic the remodeling of the ECM during OC dissemination, and in particular the degradation of the basement membrane, early-stage ECM model was developed with type I collagen topped with basement membrane proteins – type IV collagen and laminin – giving rise to an artificial basement membrane (ABM). The degraded late-stage ECM is represented here by artificial connective tissues (ACT) only made up of type I collagen (Figure 1A). These models were previously thoroughly characterized in terms of ECM structure, composition and stiffness (Figure S5 and S7) [5]. These matrices replicate the physiological stiffness range found in normal and pathological tissues, allow fluid exchange, tumor spheroid seeding and are easily integratable into microfluidic chip systems (Figure 1B-C). After spheroid seeding, fibronectin and TGF-β were added to the culture medium together or separately to model the ascitic environment in the peritoneal cavity (Figure 1D1). Fibronectin concentration in ascites varies greatly under different environments (from ∼10 μg/mL in healthy state to over 200 μg/mL in patient ascites [39]). In addition, TGF-β has been shown to increase cellular migration and decrease cellular proliferation at concentration as low as 10 ng/mL [40,41]. In this respect, the circulating medium was supplemented with fibronectin at a concentration of 50 μg/mL and TGF-β at a concentration of 10 ng/mL. Alternatively, both factors were added simultaneously, keeping each concentration identical.

### 3.1. Impact of fibronectin and/or TGF-β on OTS migration on ABM

After supplementing the culture medium with fibronectin +/-TGF-β, confocal imaging showed that similar migration patterns were observed whatever the composition of the circulating environment (Figure 2A). Qualitatively, isolated cells migrate radially out of the core part of the spheroids. Meanwhile, the spheroids maintain a densely packed core at the center. This core exhibits a low vimentin expression indicating the epithelial-like phenotype of cells in the core, compared to the vimentin-expressing cell migrating out of the spheroids. This illustrates the dynamic reorganization of the cytoskeleton, with cells transitioning towards mesenchymal-type in the course of migration.

**Figure 2.**
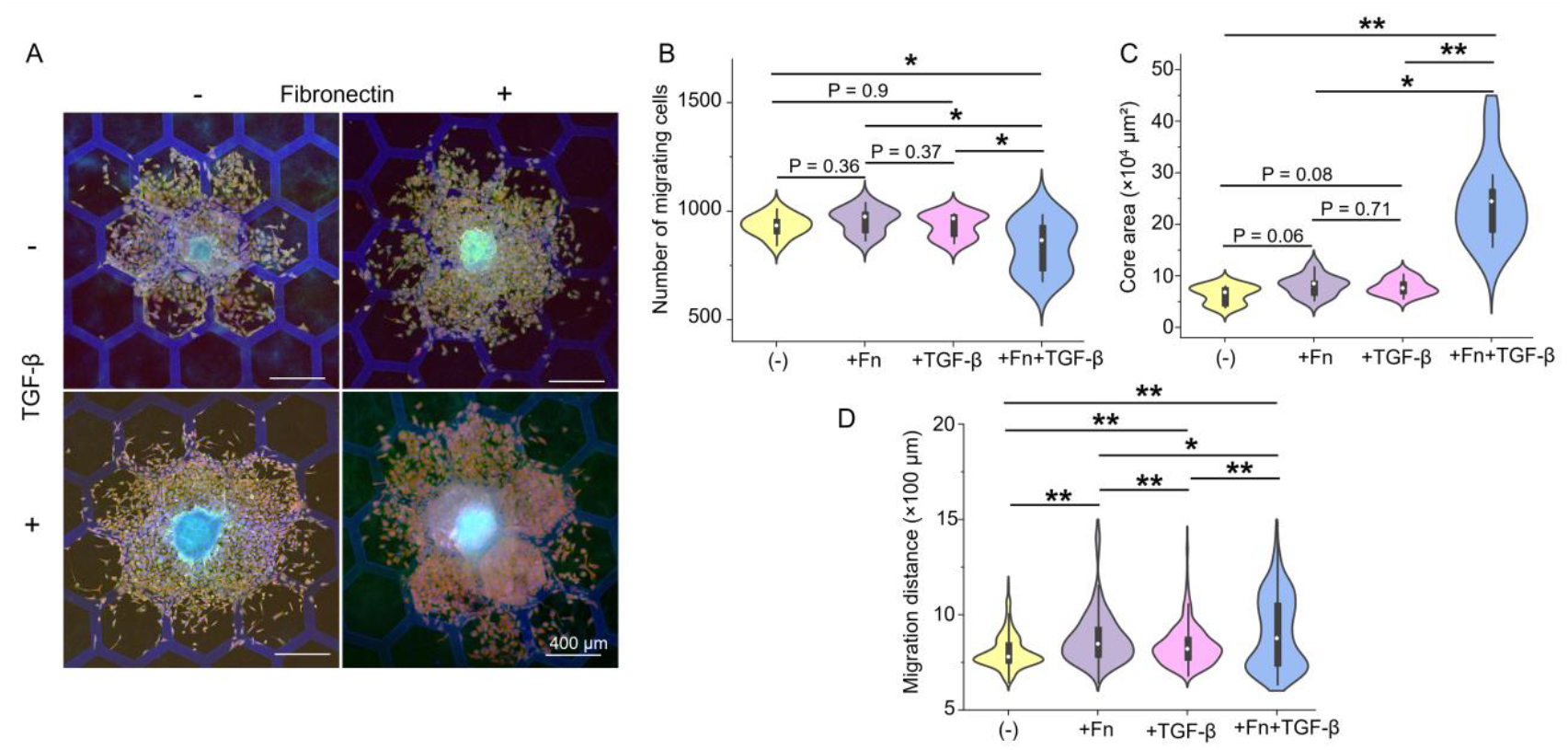
Effects of fibronectin and/or TGF-β on the migration of SKOV-3 cells on ABM. (A) IF imaging of SKOV-3 cells after 48 hours culture with fibronectin and/or TFG-β on ABM. Cells were stained for vimentin (red), actin (green), and nuclei (DAPI, blue). B-D: Plots of the number of migrating cells, core area and migration distance on ABM. Each condition using Two-Sample t-test (*p<0.05, **p < 0.01).

To quantify migration behavior, we extracted the number of isolated leading cells from nuclear staining using ImageJ and measured the area of the spheroid core, which is defined as the dense cell region where cells cannot be segmented automatically. The migration distance was calculated from the average of the 5% highest migration distances of the isolated cells, thus representing the migratory capacity of peripheral isolated cells (Figure S6).

First, in terms of number of migrating cells, no variation was observed when adding Fn or TGF-β only to the culture medium. In contrast, a significant decrease was observed in presence of both Fn + TGF-β, (836 ± 119 *vs*. culture medium (-): 934 ± 49, +Fn: 958 ± 57 and +TGF-β: 934 ± 56) (Figure 2B). Similarly, a significant increase in core area was observed when adding Fn + TGF-β, which was not observed upon addition of only one component (24×10^4^ ± 2×10^4^ μm^2^ *vs*. (-): 6×10^4^ ± 2×10^4^ μm^2^, +Fn: 8×10^4^ ± 2×10^4^ μm^2^ and +TGF-β: 8×10^4^ ± 2×10^4^ μm^2^) (Figure 2C).

The migration distance in presence of Fn or TGF-β were increased compared to the perfusion of culture medium (from 805 ± 89 μm, to +FN: 877 ± 140 μm and +TGF-β: 839 ± 109). When Fn and TGF-β were added simultaneously, an increase up to 906 ± 192 μm was measured (Figure 2D).

Interestingly, neither the fibronectin nor TGF-β alone has as strong impact on cell migration in presence of basement membrane proteins compared to the combination of Fn+TGF-β. Moreover, the impact of Fn+TGF-β was more than the addition of the impact of each component taken separately evidencing the synergistic effect of Fn and TGF-β addition.

Given this synergistic effect, the impact of ECM composition was investigated on ACT (in absence of basement membrane proteins) under perfusion of Fn+TGF-β. Under the perfusion of culture medium, OC migration on ACT shows a different pattern than on ABM with the absence of a proper spheroid core and a low vimentin expression (Figure 3A). In absence of basement membrane proteins, the number of migrating cells significantly decreases under perfusion of Fn+TGF-β (from 866 ± 71 to 646 ± 119, Figure 3B). Simultaneously, a significant increase in core area was observed from 4 ± 1×10^4^ μm^2^ to 14 ± 5×10^4^ μm^2^ (Figure 3C). Finally, no significant difference could be observed in migration distance upon perfusion of Fn+TGF-β (Figure 3D).

**Figure 3.**
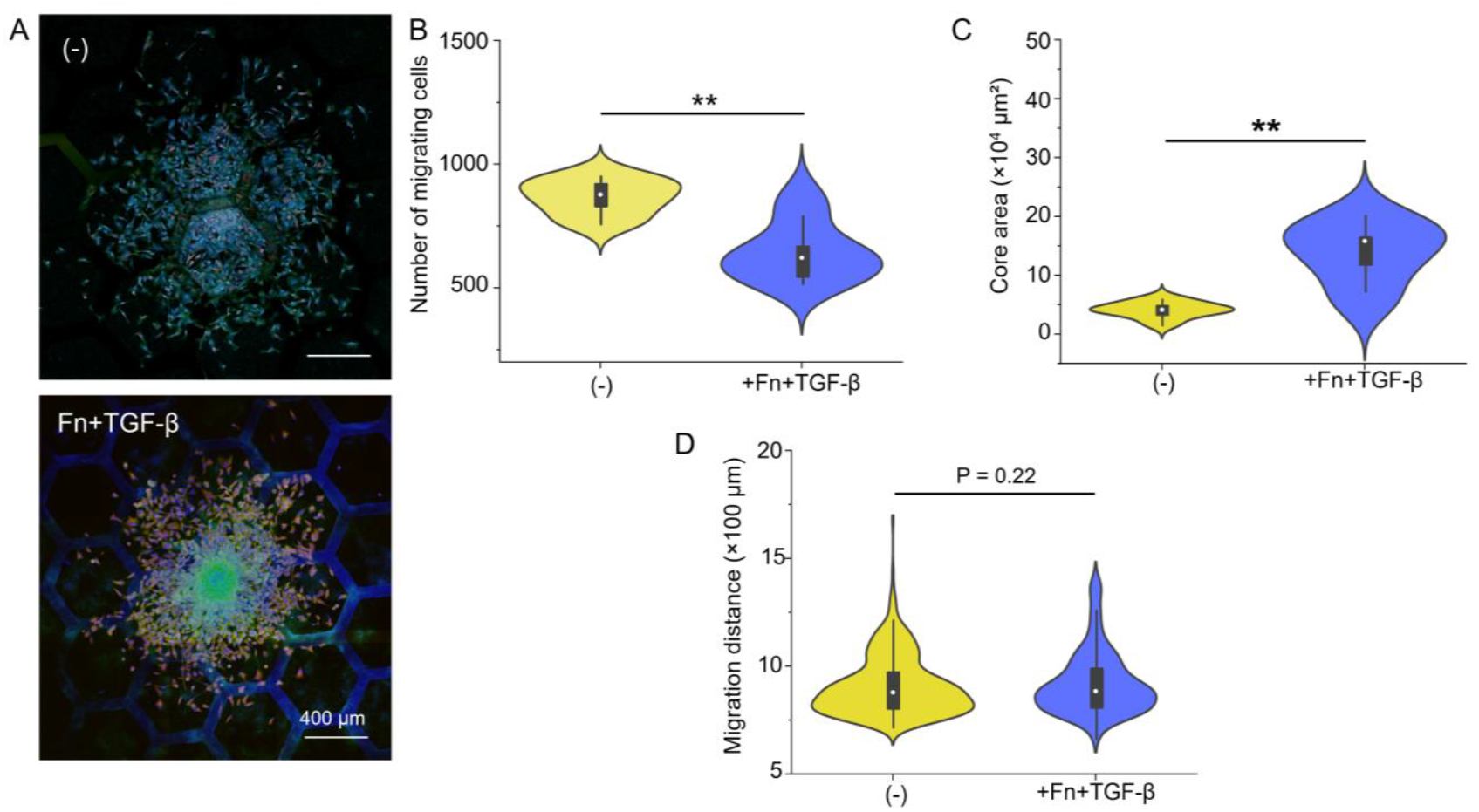
IF images of SKOV-3 cells after 48 hours culture in absence (-) or presence of fibronectin and TFG-β on ACT. Cells were stained for vimentin (red), actin (green), and nuclei (DAPI, blue). B-D. Plots of the number of migrating cells, core area and migration distance on ACT under the perfusion of medium in absence (-) or presence of Fn+TGF-β (**p < 0.01).

### 3.2. Impact of ECM and of fibronectin and TGF-β on OC cell morphology and EMT

To study the morphology of isolated cells (in the leading area) during tumor cell migration, we measured the aspect ratio and projected area of the cells and of their nuclei. We refined our analysis using Principal Component Analysis (PCA) [42]. It extracts the geometrical descriptors, or *shape modes*, that account for the morphological heterogeneity of a population of cells/nuclei [37,38]. For each descriptor, the distribution of values associated with standard deviations are calculated. In the analysis, we only accept components that account for more than 5% of variability. Cell morphology, based on the staining of the actin cytoskeleton, reveals two shape modes on ACT and ABM (Figure 4A). The most important geometrical descriptors accounting for cell morphology variations were found to be mostly the cell spreading (accounting for 75.2% of total variance) and the cell deformation, *i*.*e*., circularity (13.4% of total variance). Nuclei were also found to vary with respect to these two geometrical descriptors of spreading and deformation accounting for 77.5 and 11.9% of total variance respectively (Figure 4A).

**Figure 4.**
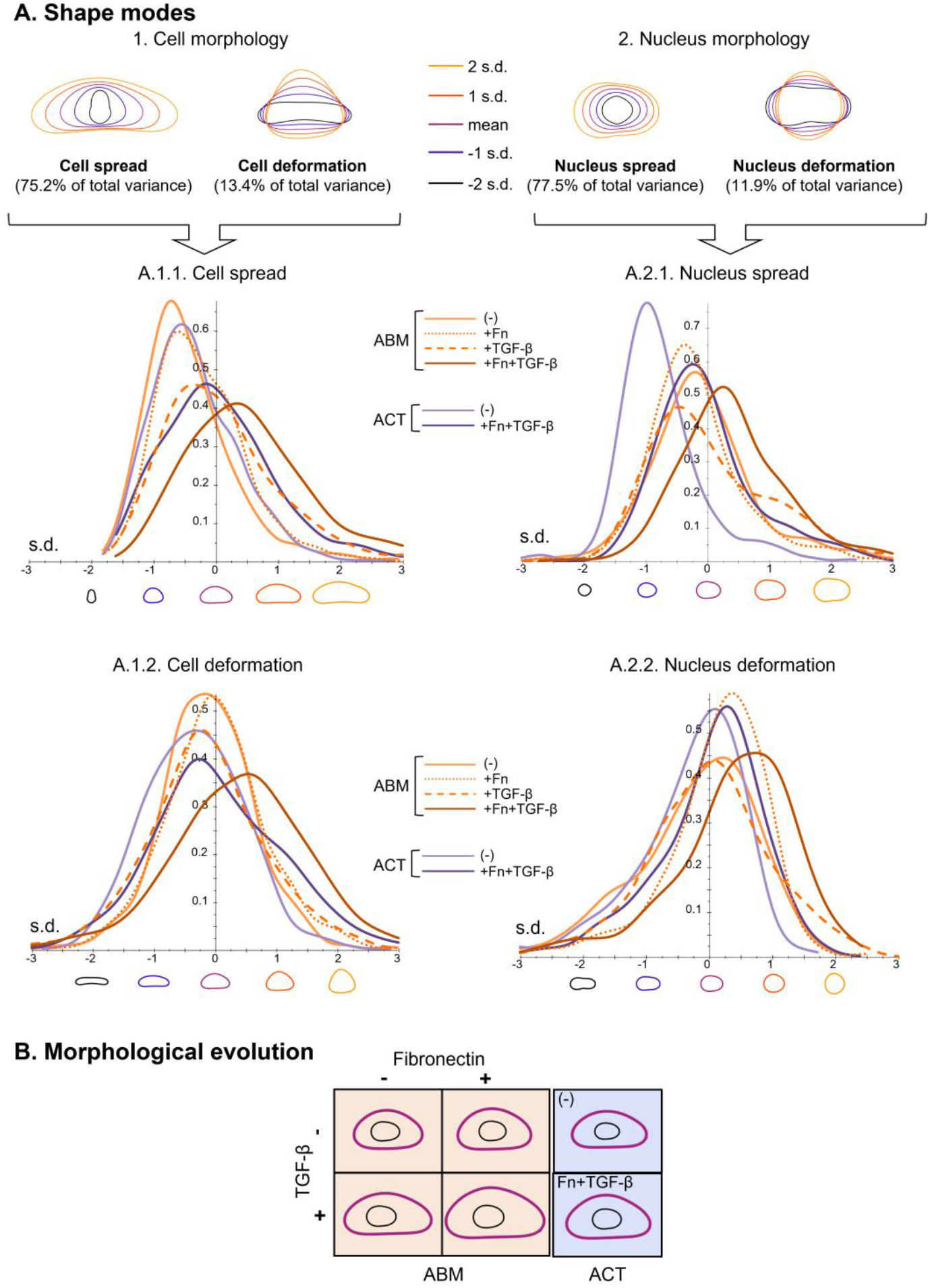
PCA of cell and nucleus morphology. (A) Shape modes describing variances in 1/ cell morphology and 2/ nucleus morphology. Cell and nucleus morphology frequency for the (A.1.1. and A.2.1.) spread and (A.1.2. and A.2.2.) deformation. (B) Comparison of the overall morphology of SKOV-3 nuclei (in black) and cells (in purple) cultured on ACT and ABM after 48 hours in different circulating environments.

#### 3.2.1. Impact of ECM and circulating environment on cellular morphology

Figure 4A also shows the cell morphology frequency. The X-axis represents the values of the principal components, with the corresponding cellular morphology schematically depicted under the X-axis. The Y-axis is the probability density. Negative values represent standard deviations below the mean. Regarding cell morphology variations as a function of circulating environment and ECM protein composition, the main geometrical descriptor – cell spread – shows a larger distribution of standard deviations, *i*.*e*., higher heterogeneity. The cell aspect ratio does not differentiate groups as clearly. In order to quantify these characteristics, we calculated medium, interquartile range (IQR) and mean value of cell spread and deformation. Considering the normal morphological distributions, IQR describes the dispersion of data around the median, *i*.*e*., quantify cell heterogeneity (Table S2).

On ABM matrices, the presence of Fn or TGF-β results in size increase, more pronounced upon addition of TGF-β, together with an increased dispersion of cell sizes (Figure 4A.1.1, Table S2). When both factors were added simultaneously, the size and dispersion increases were strikingly enhanced. In terms of cell aspect ratio, a limited shape evolution upon addition of Fn or TGF-β was observed. In contrast, Fn+TGF-β made the cells strikingly rounder, but with an increased dispersion.

Similar effects were measured on ACT: upon addition of Fn+TGF-β, an increase in median and IQR values indicated an increased cell spread and deformation with increased heterogeneity (Figure 4A.1.2, Table S2). Finally, the impact of the simultaneous addition of Fn+TGF-β was found to be much greater on ABM than on ACT.

#### 3.2.2. Impact of ECM and circulating environment on nuclear morphology

Regarding nuclear morphology, the perfusion of Fn or TGF-β on ABM induced a decrease in nucleus spread, with a greater heterogeneity in presence of TGF-β characterized by the presence of a second population with larger nuclei (Figure 4A.2.1, Table S3). In contrast, a spread increase together with a larger heterogeneity was measured when both factors were combined. A more pronounced increase on nucleus spread was observed on ACT, together with an increase heterogeneity, after addition of Fn+TGF-β (Figure 4C2, Table S3). As for the nucleus deformation, addition of Fn or/and TGF-β on ABM triggers an increase in deformation, with decreased heterogeneity. This indicates that nuclei become rounder in presence of Fn or/and TGF-β. This was also observed on ACT substrates upon perfusion of Fn+TGF-β. These results indicate that nuclei became larger and rounder upon Fn+TGF-β perfusion.

The overall morphological variations of SKOV-3 nuclei and cells are recapitulated in Figure 4B: on ABM, Fn addition to the culture medium induced an increase in the overall cell spread, whereas the addition of TGF-β not only increased cell area but also triggered distinct morphological elongation. Compared to the effects of Fn or TGF-β alone, the combined action of fibronectin and TGF-β shows a stronger trend to cell enlargement, with cells becoming much more spread than in presence of Fn or TGF-β alone. On ACT, the addition of Fn+TGF-β in the microenvironment resulted in cells becoming more spread although to a lesser extent than on ABM.

To further investigate morphological evolution of OC cells, we analyzed the reorganization of the cytoskeleton in the context of increasing cell invasiveness. To this aim, we focused on the expression of vimentin cytoskeleton protein, known to be correlated with EMT and cell invasiveness in metastatic contexts. While having a low intensity with cortical distribution in epithelial cells, mesenchymal cells exhibit a higher vimentin intensity, with cortical distribution, co-localizing with the actin cytoskeleton. Upon varying the circulating environment, we observe a significant decrease in vimentin intensity when Fn or TGF-β were perfused on ABM. Interestingly however, when Fn and TGF-β were added simultaneously, the reverse effect was observed with a large increase in vimentin intensity (Figure 5A).

**Figure 5.**
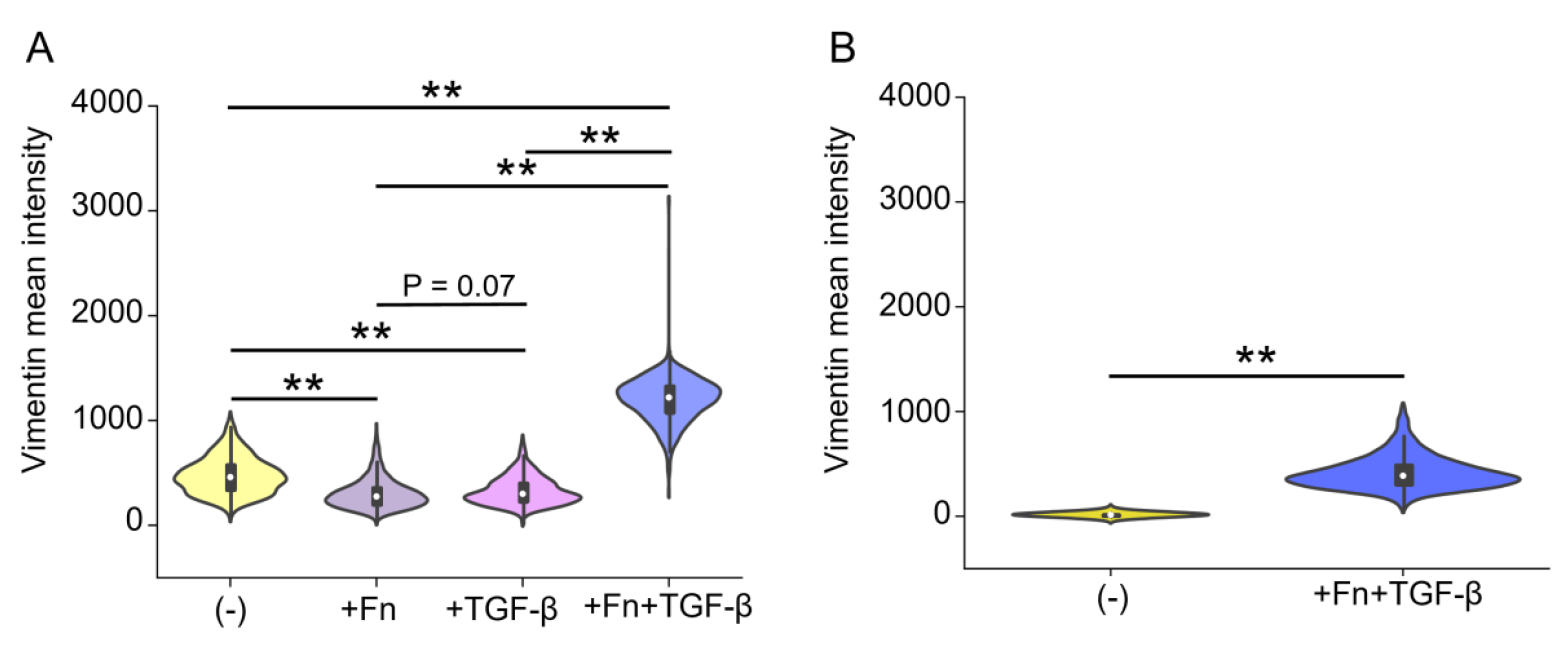
Plots of vimentin intensity as a function of the circulating environment after seeding OC cells on (A) ABM and (B) ACT (**p < 0.01).

This supports that cytoskeletal reorganization accompanying cell invasiveness requires not only TGF-β, but its combination with fibronectin. A similar large increase was also observed on ACT upon addition of Fn+TGF-β (Figure 5B). Regarding the protein composition of ECM, vimentin intensity was found to exhibit a much higher intensity on ABM than on ACT both in absence or presence of Fn+TGF-β. This indicates that ABM more effectively promotes cytoskeleton reorganization in the course of OC cells EMT compared to ACT, which corroborates our previous findings [6].

### 3.3. ECM and patient-ascites crosstalk

The obtention of patient ascites and the patient-to-patient variability makes the acquisition of triplicates difficult. To tackle this limit, two microfluidic chips were run connected in series (Figure 1D2). Given that each ECM model in the chip was seeded with four spheroids (∼1 × 10^3^ cells/spheroid), cell-secreted proteases should not significantly impact ascites composition, and the circulating volume (5 mL for 48 hours) substantially exceeded the metabolic consumption capacity of the spheroids [43]. We therefore postulated that running ascites over one chip would not significantly affect ascitic composition for the second chip. For a given ascites, one set of two connected microfluidic chips was loaded with both ACT and ABM models. This setup allowed the exact same ascites sample to simultaneously impact cells on both ACT and ABM models for a 48-hour culture time. Ascites from four different patients were perfused, each in this two-chips configuration. The migration behavior of cells in ascitic environment and as a function of ECM composition was further investigated by IF imaging, combining staining of vimentin, actin and nuclei.

Figure 6 shows one representative spheroid for each of the two patches – ABM and ACT – obtained with three ascites. One striking evidence from the representative IF imaging is the high reproducibility observed on ABM on one side, and on ACT on the other side. This concerns first the expression of vimentin, high on ABM and hardly detectable on ACT, as well as the size of the spheroid core, being much denser and smaller on ACT than on ABM. It is important to note that the ECM dependency previously observed is not only preserved, but enhanced when patient-derived ascites is perfused. Still, variations were observed when perfusing a fourth ascites (Figure S8). As deviations in cell behavior was observed for both ABM and ACT with this batch, we attributed these behaviors to ascites. Further investigations of the ascites composition would be necessary to further investigate this point and confirm this hypothesis.

**Figure 6.**
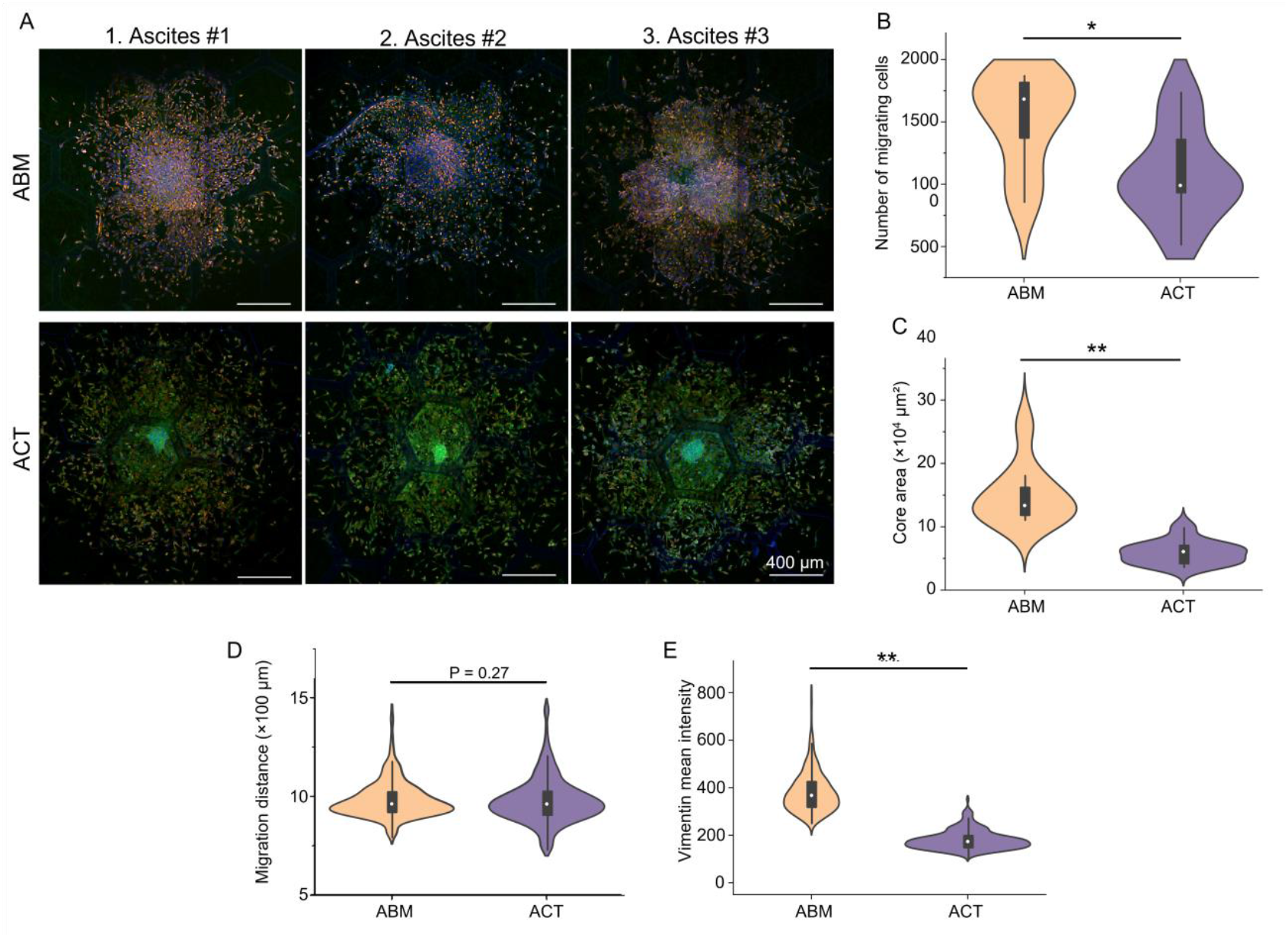
A. IF confocal imaging of SKOV-3 spheroids cultured under perfusion of patient-derived ascites for 48h. Cells were stained for vimentin (red), actin (green), and nuclei (DAPI, blue). B-E. Plots of the number of migrating cells, core area, migration distance and vimentin mean intensity of SKOV-3 cells cultured under perfusion of patient-derived ascites for 48h (*p<0.05, **p < 0.01).

The migration behavior of cells was then investigated more quantitatively. After 48 hours under patient-ascites perfusion, the number of migrating cells on ABM was found to be higher than on ACT, being 1 528 ± 371 μm on ABM *vs*. 1 094 ± 407 for ACT (Figure 6B). On the other hand, the core area decreased significantly on ACT compared to ABM, going from 15×10^4^ ± 5×10^4^ μm^2^ to 6×10^4^ ± 2×10^4^ μm^2^ for ABM and ACT respectively (Figure 6C). The presence of large cores with numerous migrating cells on ABM matches the previously described model of collective migration behavior observed in presence of type IV collagen and laminin. In contrary, on ACT, fewer isolated cells migrate out of a dense spheroid core, representative of a rather individual migration.

The migration distance (Figure 6D) did not show statistically significant difference. Finally, OC cells exhibited a significantly stronger vimentin expression on ABM than on ACT (Figure 6E). This reflects a higher degree of cytoskeleton reorganization, indicating a higher propensity in transitioning along the EMT spectrum. This suggests that ascites promotes a more invasive migration on ABM than on ACT. Finally, two-photon microscopy was used to monitor ECM invasion by SKOV-3 cells. The 3D penetration of cells could thus be evidenced into both ABM and ACT matrices (Figure S9).

## 4. Discussion

Ovarian cancer (OC) remains one of the most lethal gynecologic malignancies, primarily due to its early metastatic spread within the peritoneal cavity. The peritoneal ascites fluid and the ECM jointly shape the tumor microenvironment, yet their respective and combined impacts on OC cell behavior remain underexplored. This study presents a tumor-on-chip model to investigate how ECM composition and ascitic components regulate OC spheroid migration. By constructing two distinct ECM environments— ABM (early-stage, basement membrane-rich) and ACT (late-stage, collagen I-rich)—and perfusing them with fibronectin, TGF-β, or patient ascites under physiological shear flow, we captured key biophysical and biochemical interactions affecting tumor dissemination.

Impact of fibronectin and/or TGF-β, and of patient ascites on SKOV3 migration behavior as a function of ECM was assessed by measuring migrating cell number, core area and migration distances. Cell invasiveness was evaluated by measuring vimentin expression and cellular and nuclear morphological evolution. Figure 7A is a summary table of the evolution of these parameters upon culturing OC spheroids on ABM and ACT in different circulating environments when compared to the controlled values obtained when perfusing culture medium. Fibronectin perfusion has been shown to slightly enhance the migration distance while vimentin expression of OC cells on ABM decreased. This can be attributed to Fn role in promoting cell-to-cell adhesion, notably supporting the formation of multicellular aggregates [45], thereby favoring a collective migration. TGF-β alone had a similar effect on SKOV3 migration. Very importantly, under the combined influence of fibronectin and TGF-β, the migratory capacity of OC cells was significantly enhanced on ABM (core area, migration distance, vimentin expression). This is reminiscent of previous works that have reported that TGF-β induces EMT, further leading to fibronectin overexpression. Fibronectin, in turn, achieves a synergistic effect, promoting cancer cell adhesion and invasion [46]. Our results support the fact that the combined addition of TGF-β and fibronectin creates a pro-migratory environment that mirrors certain *in vivo* processes, enhancing OC cell migration. Interestingly, a similar behavior was observed when perfusing fibronectin and TGF-β in microfluidic chips hosting SKOV3 spheroids seeded on ACT: in absence of basement membrane proteins, larger core area and higher vimentin expression were measured under the simultaneous perfusion of Fn+TGF-β. Noticeably, a lower number of migrating cells was measured on both ABM and ACT, which supports the transition towards a more collective migration mode under the perfusion of Fn+TGF-β, where fewer isolated cells can be automatically segmented.

**Figure 7.**
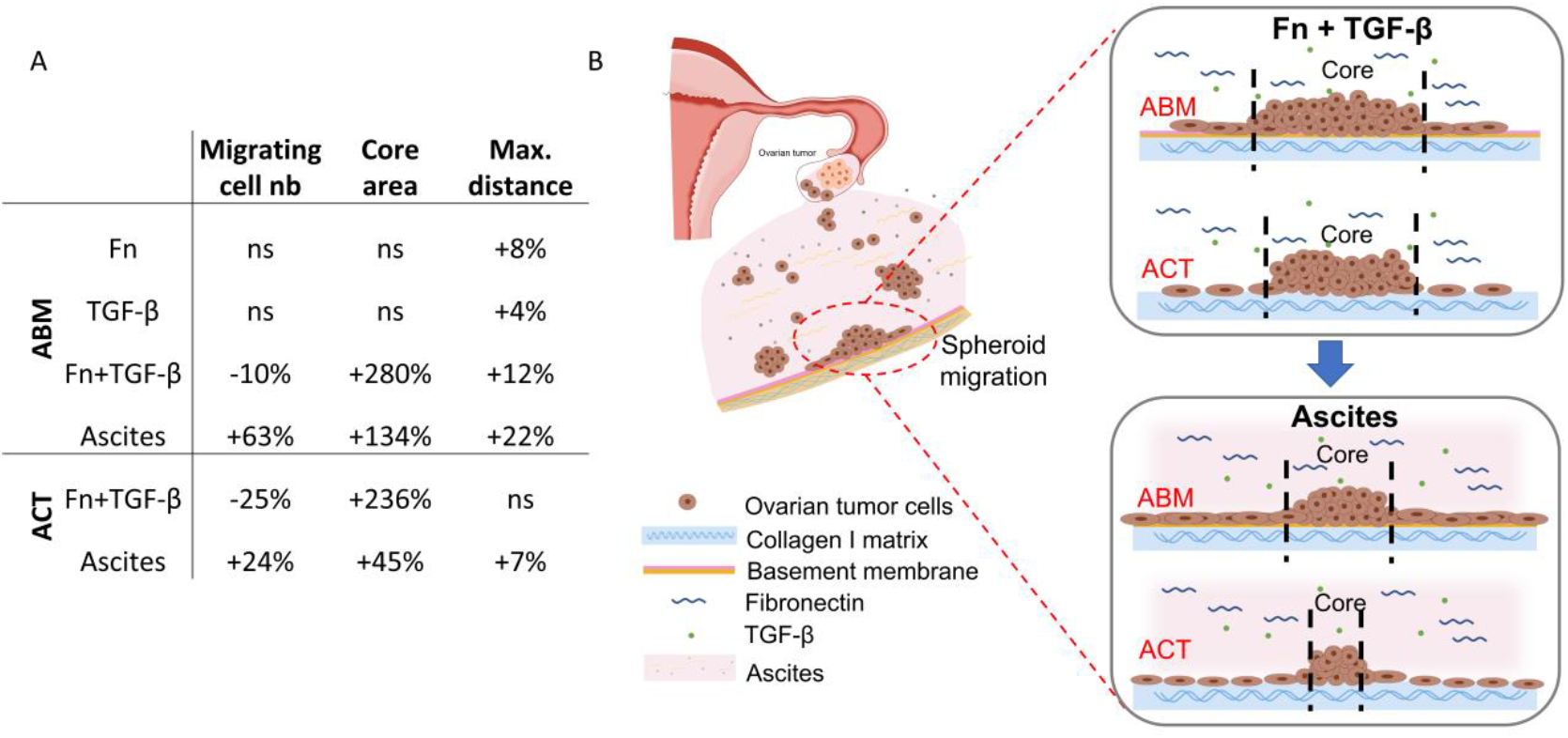
A. Evolution of migrating cell number, core area and migration distances upon varying the circulating environment and the ECM composition when compared to the perfusion of culture medium. *ns* is for non-significant. B. Schematic summarizing the impact of ECM and ascites components on OC cells migration.

Under the perfusion of ascites from patients, two key results should be highlighted: the large increase in core are and number of migrating cells, when compared to the perfusion of cell culture medium (Figure 7A). Noticeably, this corresponds to an important decrease in spheroid areas compared to the perfusion of Fn+TGF-B (Figure 7B). While this could be associated to a more individual migration mode, the associated increases in migrating cells and migration distance indicate that patient ascites significantly promotes OC cell migration on both ECM models compared to the simultaneous addition of Fn+TGF-β. More cells migrate out from the core with longer distance upon ascites perfusion than under perfusion of model ascites. Very importantly, an enhanced ECM-dependence was measured in presence of patient ascites.

The presence of the full ascites thus triggers important changes in SKOV-3 migration pattern. Meanwhile, in presence of basement membrane proteins, OC cells primarily display collective migration behavior, with a large defined core area and a higher density of isolated cells in the leading area compared to the migration behavior observed on type I collagen-rich ACT. In contrast, on the ACT model, OC cells exhibit a rather individual migration pattern, characterized by a smaller core area of the spheroids and a lower number of isolated cells in the leading area. The stiffness of ABM and ACT showed no significant difference (P > 0.05, Figure S7), which cannot be attributed the deviation in cell behavior to stiffness. Patient-ascites, rich in diverse biochemical cues, robustly promoted migration across both ECM types. However, ECM composition still defined the migration strategy. This suggests that while ascites modulates the extent of migration, ECM dictates the mode. Morphological analyses reinforced this conclusion: cells on ABM displayed greater size, roundness, and vimentin expression—hallmarks of collective and invasive migration. ACT, by contrast, promoted a more individual, mesenchymal phenotype. Our results demonstrate that ECM composition critically dictates the migration mode—collective migration on ABM *vs*. individual migration on ACT. Even under the influence of potent pro-migratory cues such as TGF-β and fibronectin, the migration pattern remained ECM-dependent. In particular, cell-ECM interactions may be strengthened by the presence of a high-density layer of collagen IV and laminin within the ABM model, which exhibits a retention capability for regulatory factors to support the stability of the cellular flow microenvironment, thereby influencing cell migration patterns.

Importantly, the tumor-on-chip platform recapitulates essential features of OC metastasis and provides a tractable system to dissect ECM–ascites–tumor interactions. This holds promise not only for mechanistic studies but also for screening anti-metastatic therapies that target ECM remodeling or tumor-stroma crosstalk.

## 5. Conclusion

This study demonstrates that the extracellular matrix (ECM) is a central determinant of ovarian cancer (OC) cell migration behavior, even within complex and clinically relevant ascitic environments. Using a physiologically relevant tumor-on-chip, we show that ECM composition—notably the presence or absence of basement membrane proteins—governs the mode of migration, with collective migration occurring on basement membrane-like ECM (ABM) and individual migration on connective tissue-like ECM (ACT).

While fibronectin and TGF-β, two key ascitic components, synergistically enhance migration and EMT features, their impact remains secondary to that of ECM composition. Perfusion of patient ascites confirmed this trend, promoting migration yet preserving ECM-dictated migration patterns.

These findings underscore the importance of incorporating ECM complexity in *in vitro* models to accurately recapitulate metastatic processes. Our work supports the development of ECM-targeted therapeutic strategies and highlights the tumor-on-chip platform as a powerful tool for dissecting tumor– microenvironment interactions and for advancing personalized approaches in ovarian cancer research.

## Author contributions

Z.W. performed the experiments, A.B. and Am.La. performed the image segmentation and the PCA analysis, L.C. and Al.Le. provided the patient ascites and helped in implementing experimentations with ascites. M.C.S.K. and C.A. performed the multiphoton experiments. C.A. designed the study and wrote the manuscript. All authors discussed the results and commented on the manuscript.

## Declaration of competing interest

The authors declare no competing interest.

## Acknowledgments

We thank the Guangzhou Elite Project for the PhD grant of Zixu Wang; Christophe Hélary and Gervaise Mosser for their help in the extraction and purification of type I collagen. This project has received financial support from the CNRS through the MITI interdisciplinary programs. Multiphoton imaging at LOB was partly supported by the Agence Nationale de la Recherche (contracts ANR-11-EQPX-0029 Morphoscope2 and ANR-10-INBS-04FBI). This work benefited from the technical contribution of the Institut Pierre-Gilles de Gennes joint service unit CNRS UAR 3750. The authors would like to thank the engineers of this unit for their advice during the development of the experiments: Bertrand Cinquin, Audric Jan, Taha Messelman. This work was funded by ANR JCJC Modulo-EMT ANR-21-CE19-0006 and is a part of Alphonse Boché doctoral work.

## Notes

### Competing Interest Statement

The authors have declared no competing interest.

