## Supplementary Information for "The extracellular matrix dictates ovarian cancer cell migration in an in vivo-derived circulating environment"

**1. Materials and methods**

***1.1. Fabrication of the porous supports***

***1.2. Spheroid growth***

***1.3. One-way self-circulation chip system***

***1.4. Simulating the flow-induced shear stress of the peritoneal cavity***

***1.5. Second harmonic generation / 2-photon excited fluorescence***

**2. ACT and ABM characterization using SEM and two photon microscopies**

**3. Quantification of spheroid migration from immunofluorescence images**

**4. Characterization of the elasticity of the ECM models as a function of protein composition, AFM nanoindentation measurements.**

**5. Image processing and statistical analysis**

***5.1. Number of isolated cells in the leading area analyzed per condition***

***5.2. Medium, interquartile range (IQR) and mean value of cell size and aspect ratio***

***5.3. Medium, interquartile range (IQR) and mean value of nucleus size and aspect ratio***

**6. Migration of SKOV-3 spheroids on ABM and ACT upon patient ascites perfusion**

**7. SKOV-3 invasion in ABM and ACT by two-photon microscopy**

### 1. Materials and methods

#### *1.1. Fabrication of the porous supports*

Porous supports similar to the ones that are commercially available from the Mesobiotech® company were fabricated by photolithography and soft-lithography in the lab. Supports were designed using CleWin5 software and printed from micropattern generator (Heidelberg µPG 101 Tabletop Micro Pattern Generator) with an array structure with a honeycomb microframe. The width of the honeycombs was set at 400 µm, with a frame width of 50 µm and a thickness of 50 µm. First, a double layer SU-8 mold was fabricated by photolithography. The mesh layer was patterned on a silicon wafer using a 50 µm thick SU-8 negative photoresist by UV exposure at 250 mJ/cm². Then, the honeycomb frame of 50 µm height was directly patterned on the mesh layer by another round of UV exposure at 250 mJ/cm². After development in propylene glycol methyl ether acetate, this double layer SU-8 mold was then exposed in trimethylchlorosilane (TMCS, Sigma, France) vapor for 10 min. Afterwards, a mixture of PDMS (GE RTV 615) pre-polymer and its crosslinker at ratio of 10:1 (w/w) was casted on the SU-8 mold. After curing at 75°C for 4 hours, the PDMS layer was peeled off. Then, this replicated PDMS structure was placed on a glass plate and a solution of a photo-crosslinking polymer (Ormostamp®, micro resist technology) was injected in the free space of the PDMS-glass assembly, followed by UV exposure at 1500 mJ/cm², and removal of the PDMS mold. The porous supports were then coated with gold by sputter deposition for improving hydrophilicity and drop casting of type I collagen by using an Emitech K675X Sputter Coater System working at 125 mA for 30 seconds.

#### *1.2. Spheroid growth*

Ovarian tumor spheroids are produced using a caged-honeycomb supports with a non-adherent coating to control their growth. First, gelatin nanofibers and agarose are coated on one side of the porous scaffold with 200 µm thickness. Second, the coated porous scaffold is maintained by a PDMS ring at the bottom of a 12-well plate (Figure S1). Finally, the SKOV-3 cell suspension is cast into the hole of the PDMS ring with cell culture medium. SKOV-3 spheroids with a stable size are obtained after 3 days of culture.

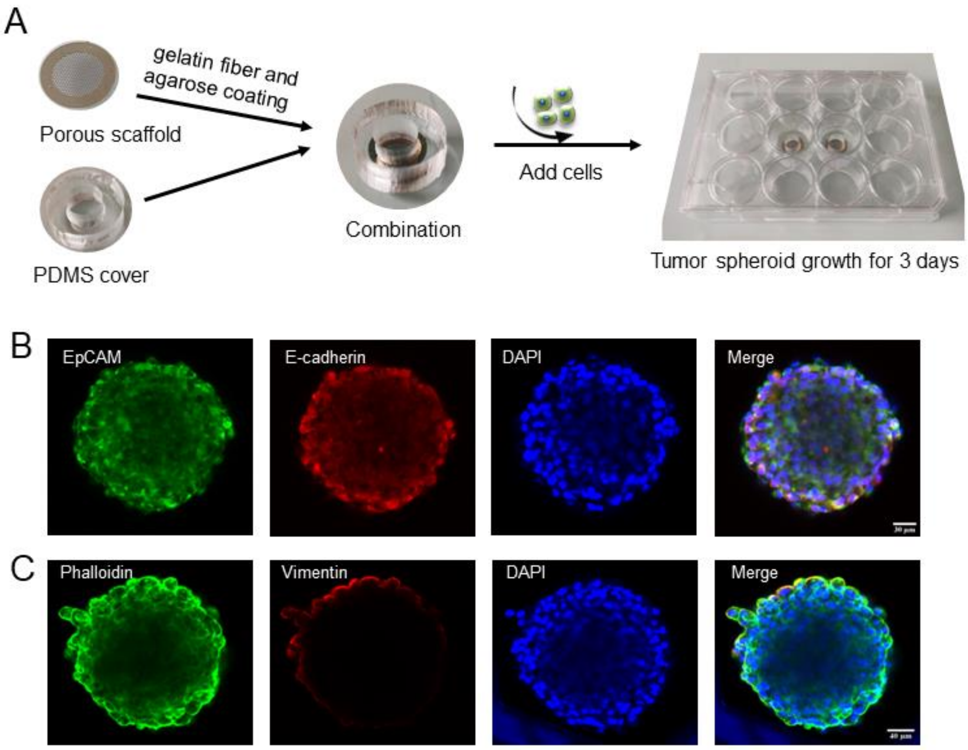


**Figure S1.** (A) Scheme of the preparation of spheroids. IF confocal imaging of a SKOV3 tumor sphere stained (B) epithelial (EpCAM and E-cadherin), and (C) mesenchymal markers (vimentin). The scale bar is 40 µm.

Figure S2 shows the resulting SKOV-3 spheroids with the uniform size of about 200 µm. The obtained spheroids can be extracted and seeded one by one. Immunofluorescence (IF) of the resulting spheroids confirms their 3D organization, as well as the presence of the epithelial markers EpCAM and E-cadherin. In contrast, staining of the vimentin cytoskeleton protein, which is commonly used as a mesenchymal marker could not be identified (Figure S2). After seeding, tumor spheroids were incubated on the ECM models for 2 hours before their transfer in the chip for culturing under perfusion for two days.

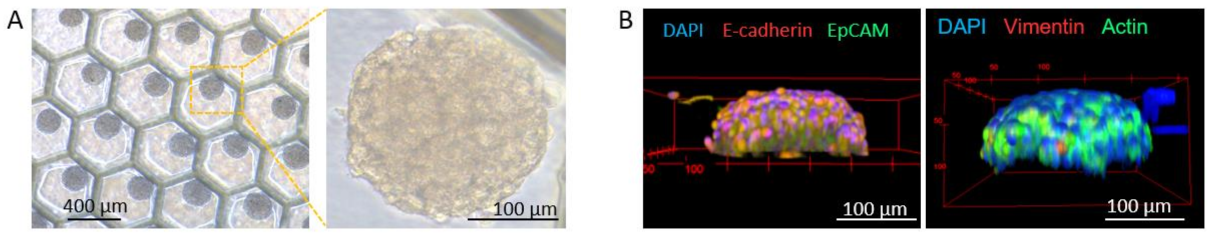


**Figure S2.** (A) Optical microscopy and (B) IF images of spheroids grown in deep-well patches. Cells are stained for DAPI, E-cadherin, EpCAM (left image), and for DAPI, vimentin and actin (right image).

##### *1.3. One-way self-circulation chip system*

We designed a one-way self-circulation chip system to cope with the limits associated with syringe pump-driven microfluidic systems: 1/ operations caused by the inability to circulate for a long time; 2/ leakage occurrence due to high pressure in the chip, while ensuring that the culture medium flows in only one direction inside the chip during the experiment.

To solve these problems, various improvements have been implemented (Figure S3):

1. Switch from *injection* to *withdraw* mode: The medium flows out of the collector through the inlet and into the chip, and then flows out of the chip from the outlet and into the syringe. The one-way valve of the injection pipe prevents air/medium from flowing into the syringe. This reduces unnecessary pressure increase in the chip

2. Use of a *one-way valve* for injection. The culture medium enters the collector through the directly connected tube. The one-way valve of the withdraw pipe prevents the culture medium from flowing back into the chip.

3. *Circulation*. After programming the pump with repeats of withdraw and injection steps, we can control the overall culturing time.

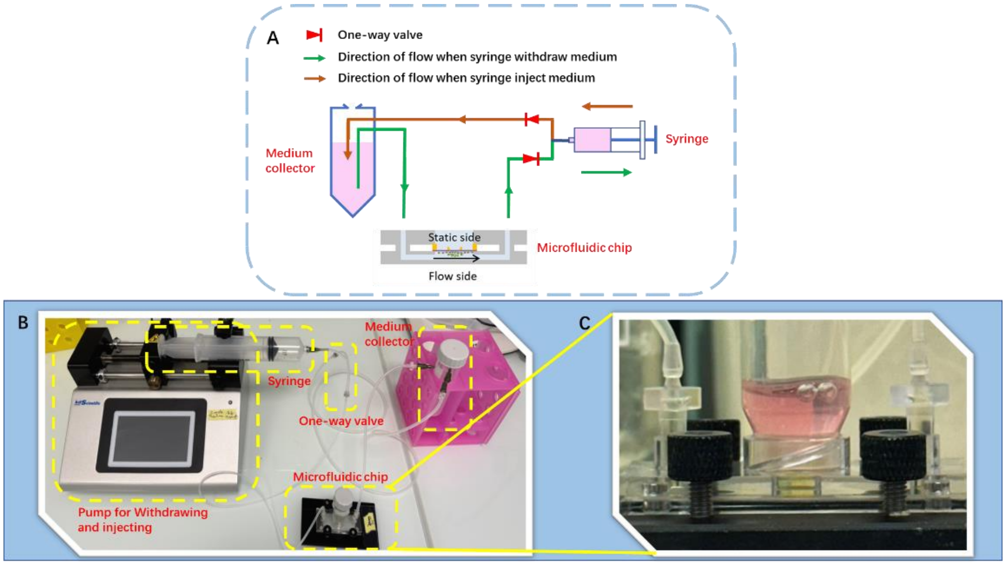


**Figure S3.** A. Schematic diagram of the one-way self-circulation chip system. B. Photo of the self-circulation system powered by a syringe pump. C. Working status of chip perfusion of patient-derived ascites.

##### *1.4. Simulating the flow-induced shear stress of the peritoneal cavity*

Based on similar setups in the literature, we chose our flow to be around some tens of µL/min, as it matches with a physiologically relevant shear stress less than a dyn.cm$^{-2}$ for our setup. Besides, with our values, our flow was laminar according to (Eq.1).

$$Re = \frac{u_{(cm/s)} * L_{(cm)} * \rho_{(kg/m^3)}}{10 * \mu_{(dyn.s/cm^2)}} < 2000 \qquad \text{(Eq.1)}$$

where µ is the dynamic viscosity of water ($\mu \approx 0.01$ dyn.s/cm² at 20°C), L the typical dimension of the channel (L = 0.5 mm) and ρ the density of water ($\rho = 998$ kg/m$^3$ at 20°C). Consequently, computer fluid dynamics modeling for laminar flow was used. We assumed our fluid to be incompressible and Newtonian, and our system to be time independent. The structure of our chip was built with rectangular channels of width and height 500 µm, and central cavity of radius 4 mm and height 900 µm. The patch was superimposed to the central cavity and was 200 µm high. The inlet channel, which goes along the x axis (with z axis being defined orthogonally to the chip) is divided into six smaller channels to spill the flow into the central cavity while reducing the shear stress. As simulated by computer fluid dynamics modeling, this allows the shear stress due to the fluid to be homogenized within the chip. Because of the symmetry properties of our chip, only half of the chip was modeled to ease computations, with a condition of symmetry in the plan y = 0. Other boundaries were considered as no-slip walls, except from the inlet and outlet (the difference in pressure was defined as pressure imposed by the syringe actuator minus ambient pressure). The patch was approximated by a porous layer of collagen I, while the chip was made of PDMS, and filled with water. A single-phase laminar flow model was used to describe the flow, coupled with a Darcy's law in the porous area at the top of the central cavity. Porosity and viscosity required for the simulation were determined from material. A physics-controlled mesh with fine element size and without further modifications was used. Results were analyzed by plotting iso-pressure lines as well as velocity and shear stress mapping with different initial pressure (Figure S4).

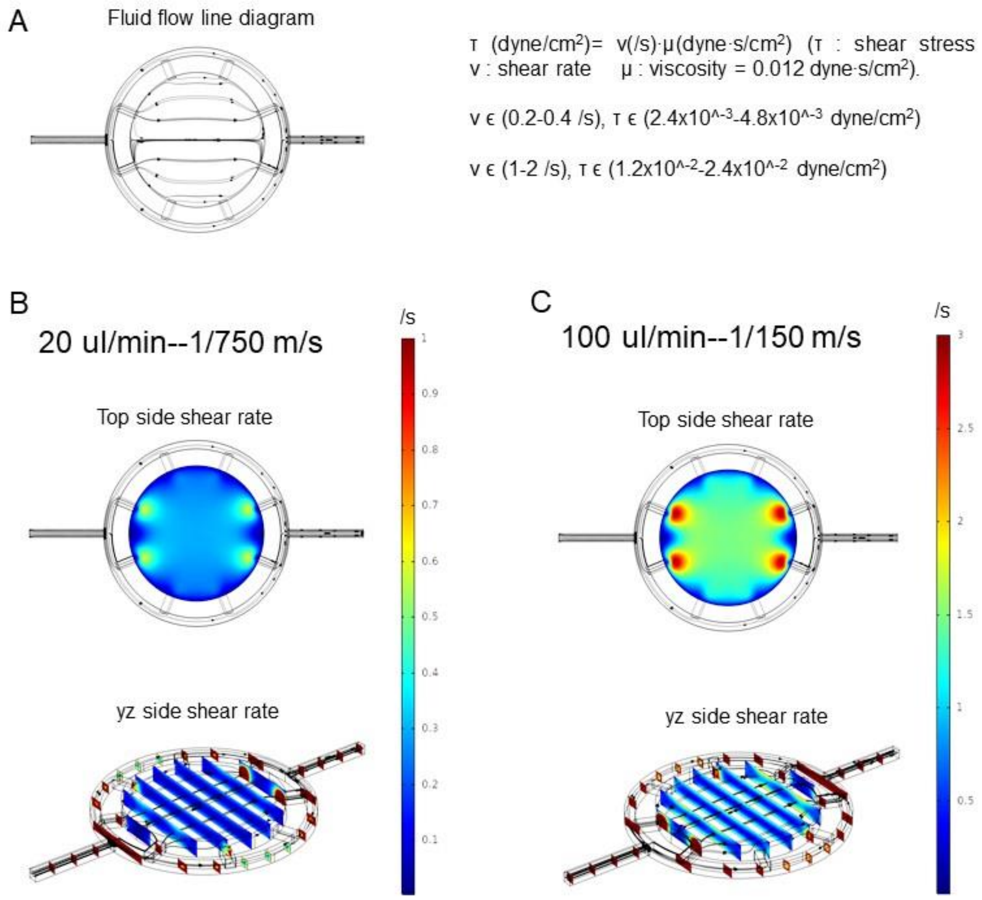


**Figure S4.** (A) Simulation of fluid flow direction and distribution in circular microchamber using Comsol Multiphysics. (B,C) Distribution of the top wall shear stress in the chamber for flow rates of 20 and 100 µl.min$^{-1}$.

##### *1.5. Second harmonic generation / 2-photon excited fluorescence*

We used a custom-built laser-scanning multiphoton microscope and recorded second harmonic generation (SHG) and 2-photon excited fluorescence (2PEF) images in parallel as previously described.39 Excitation was provided by a femtosecond titanium–sapphire laser (Mai-Tai, Spectra-Physics) tuned to 860 nm, scanned in the XY directions using galvanometric mirrors and focused using a 25× with 1.05 NA objective lens (XLPLN25X-WMP2, Olympus), with a resolution of 0.35 µm (lateral) × 1.2 µm (axial) and a Z-step of 0.5 µm for the acquisition of Z-stack images. We used circular polarization in order to image all structures independently of their orientation in the image plane, using 100 kHz acquisition rate, 420×420 nm² pixel size and a typical power of 22 mW.

For the characterization of the basement membrane by 2PEF (Figure S5), ECM models were stained with Alexa Fluor 488®-conjugated collagen IV monoclonal antibody (eBioscience™, 53-9871-82) incubated overnight at 4°C or 2 h at room temperature at a concentration of 10 µg.mL-1, together with laminin polyclonal primary antibody (Thermo Fisher Scientific, PA1-16730) overnight at 4°C or 2 h at room temperature at a concentration of 1 µg.ml-1, followed by 1 to 2 h incubation at room temperature with a goat–anti rabbit-Alexa Fluor 610 secondary antibody at a dilution of 1/250.

#### 2. ACT and ABM characterization using SEM and two photon microscopies

For SEM characterization, samples were dried at 60°C overnight, then coated with gold for 60 s using an Emitech K675X Sputter Coater System with a sputtering current of 50 mA before imaging. Samples were fixed on conductive-tapes for imaging with a TM3030 Tabletop Microscope (Hitachi High-Technologies Corporation, Japan) working at an acceleration voltage of 15 kV.

Figure S1 shows that the collagen I layer is homogeneous, with a thickness of a couple of microns. After adding successively type IV collagen and laminin, the layer thickness and aspect did not change under SEM (Figure S5B1). Combining SHG with 2-photon excited fluorescence (2PEF) allows to specifically identify the presence of each protein: a top layer of laminin, with type IV collagen embedded within the ABM (Figure S5B2).

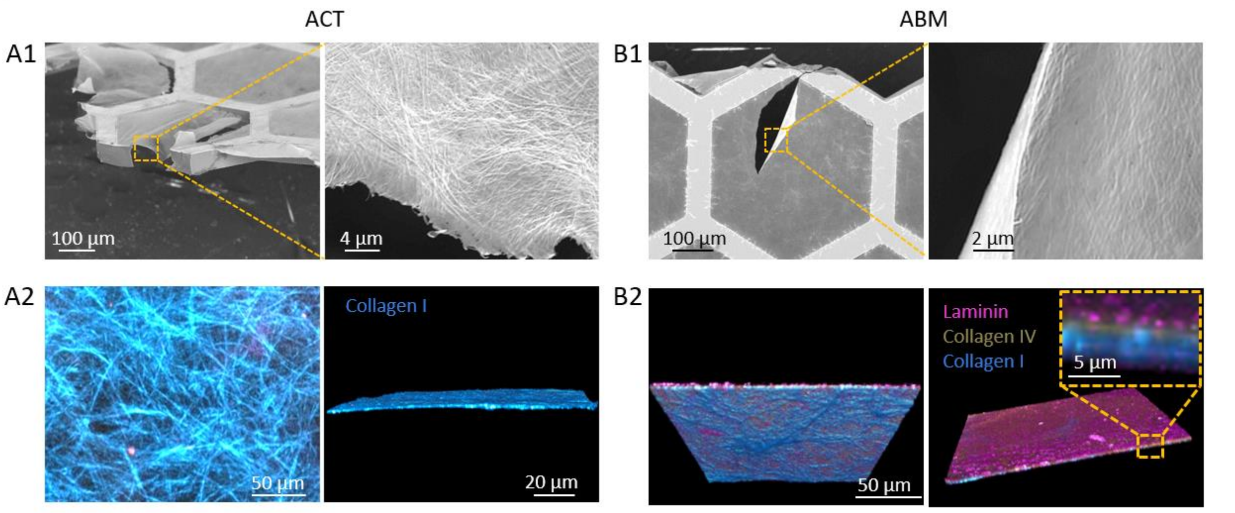


**Figure S5.** (A1,B1) SEM and (A2,B2) SHG/2-PEF images of (A) ACT and (B) ABM on honeycomb frame supports.

#### 3. Quantification of spheroid migration from immunofluorescence images

To quantify migration behavior, we measured the number of migrating cells, the area of the spheroid core and the maximum migration distance after SKOV-3 spheroid seeding on ECM models. The "core area" is defined as the dense cell region where cells cannot be segmented automatically (Figure S6A). The migration distance was measured from nuclear staining using ImageJ and extracted by averaging the 5% highest migration distances of the isolated cells (Figure S6B).

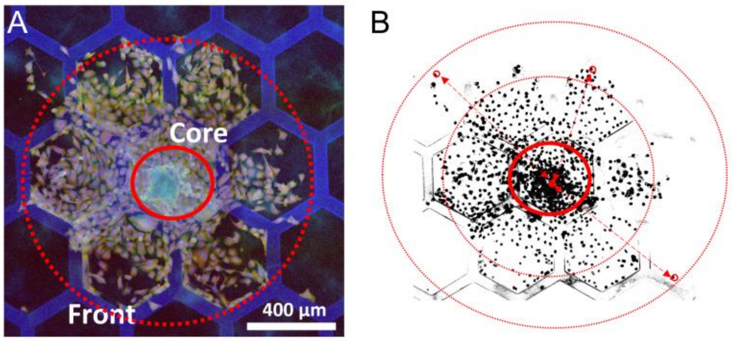


**Figure S6.** A. Definition of core part and front part of spheroids. (B) Quantification of the migration distance from the center of the core part by the segmentation of nuclei.

#### 4. Characterization of the elasticity of the ECM models as a function of protein composition, AFM nanoindentation measurements

AFM experiments were performed on hydrated ECM-models in cell culture medium at room temperature using a NanoWizard 4 (JPK BioAFM, Berlin, Germany) mounted on an Axio Observer microscope (Zeiss, Oberkochen, Germany) placed on a vibration isolation table. Before each experiment, the cantilever spring constant was accurately determined upon calibration in cell culture medium by the thermal noise method.

Young's moduli were determined by colloidal probe force spectroscopy using a gold coated cantilever (0.01 N/m) equipped with a 6.44 µm bead probe (NanoAndMore, Paris, France). Approach and retraction speeds were kept constant at 5 µm/s, ramping the cantilever by 10 µm with a 0.4 nN threshold in a closed z loop. In this option, the feedback system readjusts the initial piezo-position for each force–displacement ramp so that the maximal force applied to the sample remains constant.

AFM force-distance curves were transformed to force-indentation curves and fitted using the JPK data processing software. Curves were fitted using the contact-point independent linear Hertz-Sneddon model. The Hertz model assumes infinite sample thickness, which was approximated by using small indentation (typical indentation depth 500 nm). Automated curve fitting was applied using fitting range of 100% of curve. Force-distance curves were measured in at least 8 positions for 3 different areas of each ECM model.

Statistical analysis was performed using Origin software. Analysis of variance (ANOVA) with the Tukey's Multiple Comparison test was used for all multiple group experiments. P values < 0.05 were deemed significant. Values in graphs are the mean and standard error of mean ( $*p < 0.05$, $**p < 0.001$, $***p < 0.0001$).

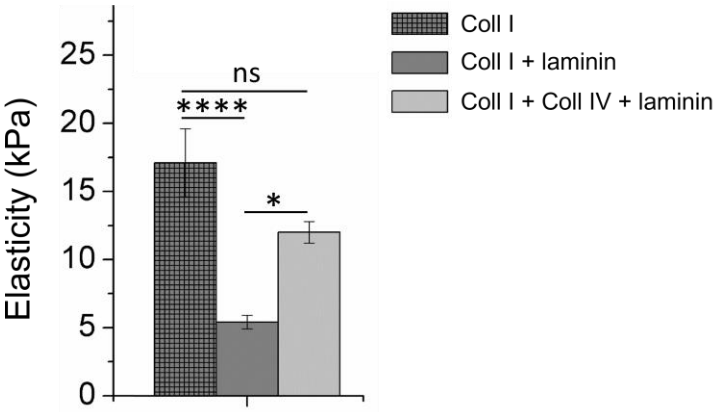


**Figure S7.** Mean Young's modulus values for all ECM models measured by peak force tapping AFM-nanoindentation. In the plots, the column represents Mean with SEM. (∗p < 0.05, ∗∗∗p < 0.00001, ns for non-significant; using Tukey's Multiple Comparison test).

#### 5. Image processing and statistical analysis

##### *5.1. Number of isolated cells in the leading area analyzed per condition*

| ECM model | Perfusion conditions | Number of cells |
|---|---|---|
| ACT | Culture Medium (CM) | 569 |
| ACT | CM + Fn + TGF-β | 678 |
| ABM | CM | 1107 |
| ABM | CM + Fn | 1006 |
| ABM | CM + TGF-β | 1153 |
| ABM | CM + Fn + TGF-β | 1020 |

**Table S1.** Overall number of isolated cells analyzed in the leading area per condition.

##### *5.2. Medium, interquartile range (IQR) and mean value of cell spread and deformation.*

| | Cell spread | | | Cell deformation | | |
|---|---|---|---|---|---|---|
| | **Median** | **IQR** | **Mean** | **Medium** | **IQR** | **Mean** |
| ABM CM | -0.57* | 0.84 | -0.42 | -0.15 | 1.00 | -0.11 |
| ABM Fn | -0.36 | 0.90 | -0.23 | -0.06 | 1.02 | 0.018 |
| ABM TGF-β | -0.09 | 1.14 | 0.17 | -0.18 | 1.14 | -0.15 |
| ABM Fn + TGF-β | 0.36 | 1.26 | 0.62 | 0.42 | 1.38 | 0.41 |
| ACT CM | -0.45 | 0.90 | -0.33 | -0.36 | 1.08 | -0.32 |
| ACT Fn + TGF-β | -0.06 | 1.14 | 0.077 | -0.12 | 1.38 | 0.049 |

**Table S2.** Median, IQR and mean values of cell spread and deformation standard deviations. Negative values represent standard deviations below the mean.

***5.3. Medium, interquartile range (IQR) and mean value of nucleus spread and deformation.***

| | **Nucleus spread** | | | **Nucleus deformation** | | |
|---|---|---|---|---|---|---|
| | **Median** | **IQR** | **Mean** | **Medium** | **IQR** | **Mean** |
| ABM CM | -0.18 | 1.00 | -0.053 | -0.09 | 1.30 | -0.21 |
| ABM Fn | -0.33 | 0.84 | -0.18 | 0.24 | 0.90 | 0.11 |
| ABM TGF-β | -0.21 | 1.32 | 0.15 | 0.00 | 1.2 | -0.043 |
| ABM Fn + TGF-β | 0.30 | 1.02 | 0.47 | 0.51 | 1.08 | 0.36 |
| ACT CM | -0.87 | 0.72 | -0.75 | -0.09 | 0.96 | -0.31 |
| ACT Fn + TGF-β | 0.15 | 0.90 | 0.0093 | 0.06 | 0.96 | -0.013 |

**Table S3.** Median values, IQR and mean values of nucleus spread and deformation standard deviations.

#### 6. Migration of SKOV-3 spheroids on ABM and ACT upon patient ascites perfusion

Figure S8 shows IF images representative of spheroid cultured on ABM and ACT under patient ascites perfusion. Noticeably, high reproducibility is observed regarding SKOV-3 migration mode on ABM on one side, and on ACT on the other side. Reproducibility concerns first the expression of vimentin, high on ABM and hardly detectable on ACT, as well as the size of the spheroid core, being much denser and smaller on ACT than on ABM. This homogeneity of cell behavior from one ascites to another one was rather not evident at first, considering the high variability of ascites composition from one patient to another one. However, variations were observed when perfusing ascites #4. As deviations in cell behavior was observed for both ABM and ACT models with this particular batch, we attributed these behaviors to ascites. Further investigations of the ascites composition would have been necessary to further investigate this point and confirm this hypothesis.

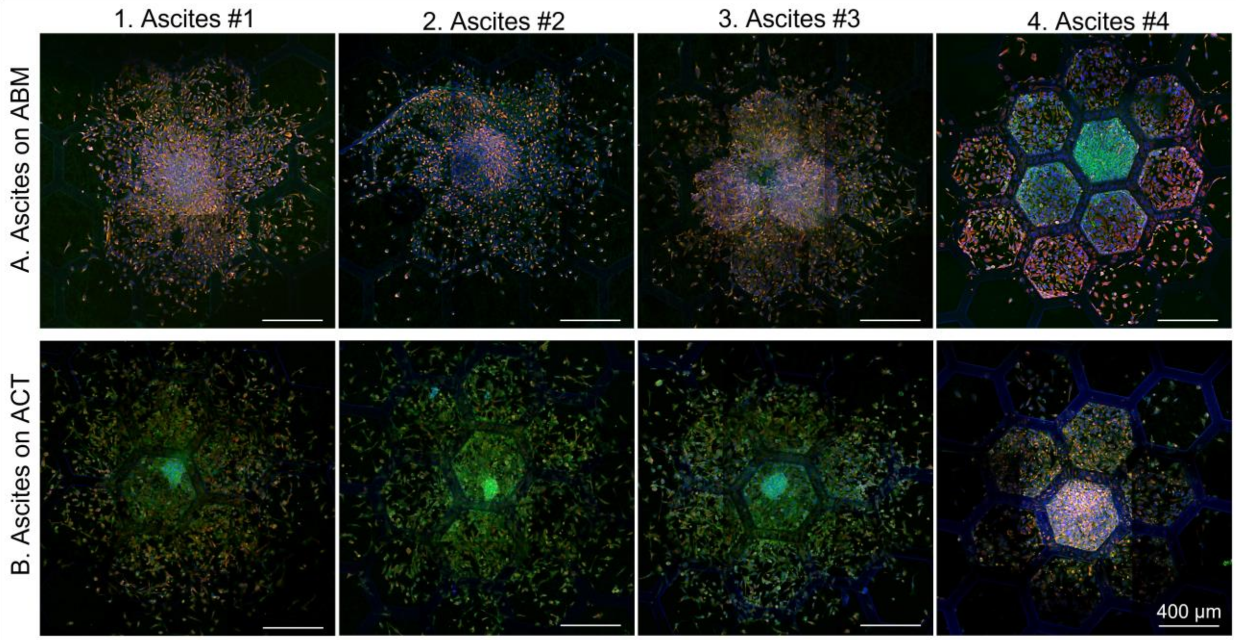


**Figure S8.** IF confocal imaging of SKOV-3 spheroids cultured under perfusion of patient-derived ascites for 48h. Cells were stained for vimentin (red), actin (green), and nuclei (DAPI, blue).

#### 7. SKOV-3 invasion in ABM and ACT by two-photon microscopy

The interactions of cells with the fibrillar collagen of the matrix were further monitored using two-photon microscopy. This allows to combine the fluorescence imaging of cells by two photon-excited fluorescence (2PEF), and the endogenous SHG signal, which is highly specific for densely packed triple-helical structures in fibrillar collagen. After perfusing ascites for 48 hours, SKOV3 spheroids were thus fixed and stained for vimentin.

As shown in Figure S9, SKOV-3 cells on both ABM and ACT matrices exhibit invasive behavior with a 3D spheroid core and migrating cells at the edge of the spheroid becoming more elongated. The side view of the 3D reconstruction also shows that migrating cells invade the collagen matrices: despite the thinness of the ECM matrix, cells invade the collagen layer, with (ABM) or without (ACT) basement membrane proteins (note that type IV collagen and laminin in the ABM model are not stained) (Figure S9A4, B4 and zoom-in in inserts). This reflects the potential of our model to investigate tumor invasion considering the three-dimensional topological structure of the matrix microenvironment. In future, we will focus more on three-dimensional matrix models.

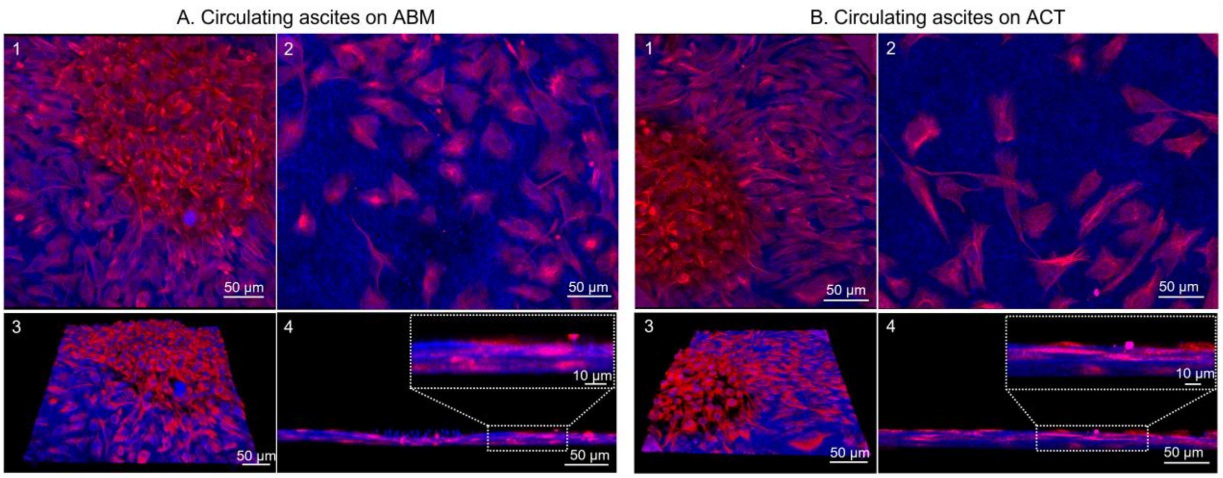


**Figure S9.** SHG/2-PEF images of core and edge area of SKOV3 spheroids seeded on (A) ABM and (B) ACT for 48 hours under perfusion of patient derived-ascites. Cells were stained for vimentin (red). (1,3) Show the core of the spheroid and (2,4) show the spheroid edge with (1,2) intensity projection and (3,4) 3D reconstruction. The inserts in A4 and B4 highlight the 3D invasion of cells within both ABM and ACT matrices.
